# A study of the structural properties of sites modified by the *O*-linked 6-N-acetylglucosamine transferase

**DOI:** 10.1101/115121

**Authors:** Thiago Britto-Borges, Geoffrey J. Barton

**Affiliations:** Division of Computational Biology, School of Life Sciences, University of Dundee, Dow Street, Dundee, DD1 5EH, UK'

**Keywords:** post-translational modification, computational biology, structural analysis, O-GlcNAc, protein disorder

## Abstract

Protein O-GlcNAcylation (O-GlcNAc) is an essential post-translational modification (PTM) in higher eukaryotes. The O-linked β-N-acetylglucosamine transferase (OGT), targets specific Serines and Threonines (S/T) in intracellular proteins. However, unlike phosphorylation, fewer than 25% of known O-GlcNAc sites match a clear sequence pattern. Accordingly, the three dimensional structures of O-GlcNAc sites were characterised to investigate the role of structure in molecular recognition. Of the 143/1,584 O-GlcNAc sites in 620 proteins were mapped to protein X-ray structures. The modified S/T were 1.7x more likely to be annotated in the REM465 field which defines missing residues in a protein structure, while 7 O-GlcNAc sites were solvent inaccessible and unlikely to be targeted by OGT. The 132/143 sites with complete backbone atoms clustered into 10 groups, but these were indistinguishable from clusters from unmodified S/T. This suggests there is no prevalent three-dimensional motif for OGT recognition. Predicted features from the 620 proteins were compared to unmodified S/T in O-GlcNAcylated proteins and globular proteins. The Jpred4 predicted secondary structure shows that modified S/T were more likely to be coils. 5/6 methods to predict intrinsic disorder indicated O-GlcNAcylated S/T to be significantly more disordered than unmodified S/T. Although the analysis did not find a pattern in the site three-dimensional structure, it revealed the residues around the modification site are likely to be disordered and suggests a potential role of secondary structure elements in OGT site recognition.

## I. Introduction

Protein *O*-GlcNAcylation, or *O*-GlcNAc, is a dynamic, intracellular glycosylation essential to mammalian development^1,2^. In animals, two enzymes mediate this post-translational modification: the glycosyltransferase *O*-linked 6-N-acetylglucosamine transferase (OGT), which adds a single, non-extensible *O*-GlcNAc moiety to serine/threonine (S/T) in the target protein; and the hexosaminidase *O*-GlcNAcase (OGA) that removes it. UDP-GlcNAc, the sugar donor to the protein *O*-GlcNAcylation, is a product of the hexosamine pathway, hence the concentration of intracellular glucose and the degree of protein *O*-GlcNAcylation levels are associated^3, 4^. At the physiological level, dysfunction of OGT activity has been linked to disease of the cardiovascular system, diabetes, impaired development, cancer and neurodegeneration^5–9^. At the cellular level, protein *O*-GlcNAcylation acts with phosphorylation, ubiquitylation and other reversible post-translational modifications in a network of cell signalling events that promote cellular adaptation to the viral infection process^10^, regulation of transcription^11^ and metabolism^12,13^.

Technical advances in mass spectrometry have led to an increase in the number of experimentally determined *O*-GlcNAc sites from 50 in the year 2000 to more than 1,000 today^14^. However, there are still obstacles to mapping *O*-GlcNAc sites reliably. The modification has a low abundance^15^ and is ten times less common than protein phosphorylation^16^. Thus, the unmodified version of the peptide can suppress the *O*-GlcNAcylated peptide mass/charge signal. In addition, methods to enrich *O*-GlcNAcylated peptides in samples have limited specificity^16, 17^, and the β-glycosidic bond is labile under the peptide fragmentation step which determines the modification's position within the peptide fragment.

Two machine learning methods have been used to detect patterns in the sequence of *O*-GlcNAc sites with limited success^20^. One of the limiting factors was that, unlike phosphorylation sites, *O*-GlcNAc sites lack a clear pattern in the primary structure. This is illustrated in Figure 1 which compares the relative sequence entropy for sites modified by OGT and three protein kinases in the PhosphoSitePlus database^14^. The observed relative entropy for OGT sites shows small signal, in contrast to protein kinase A (PKA; peak in −3 and +2), protein kinase C (PKC; peak in −3) and casein kinase 2 (CK2, peak in +3) sites. This implies that the sequence in the sites recognised by OGT carries less information than those recognised by PKA, PKS or CK2 and so are harder to distinguish from unmodified sites by sequence alone.

**Figure 1:**
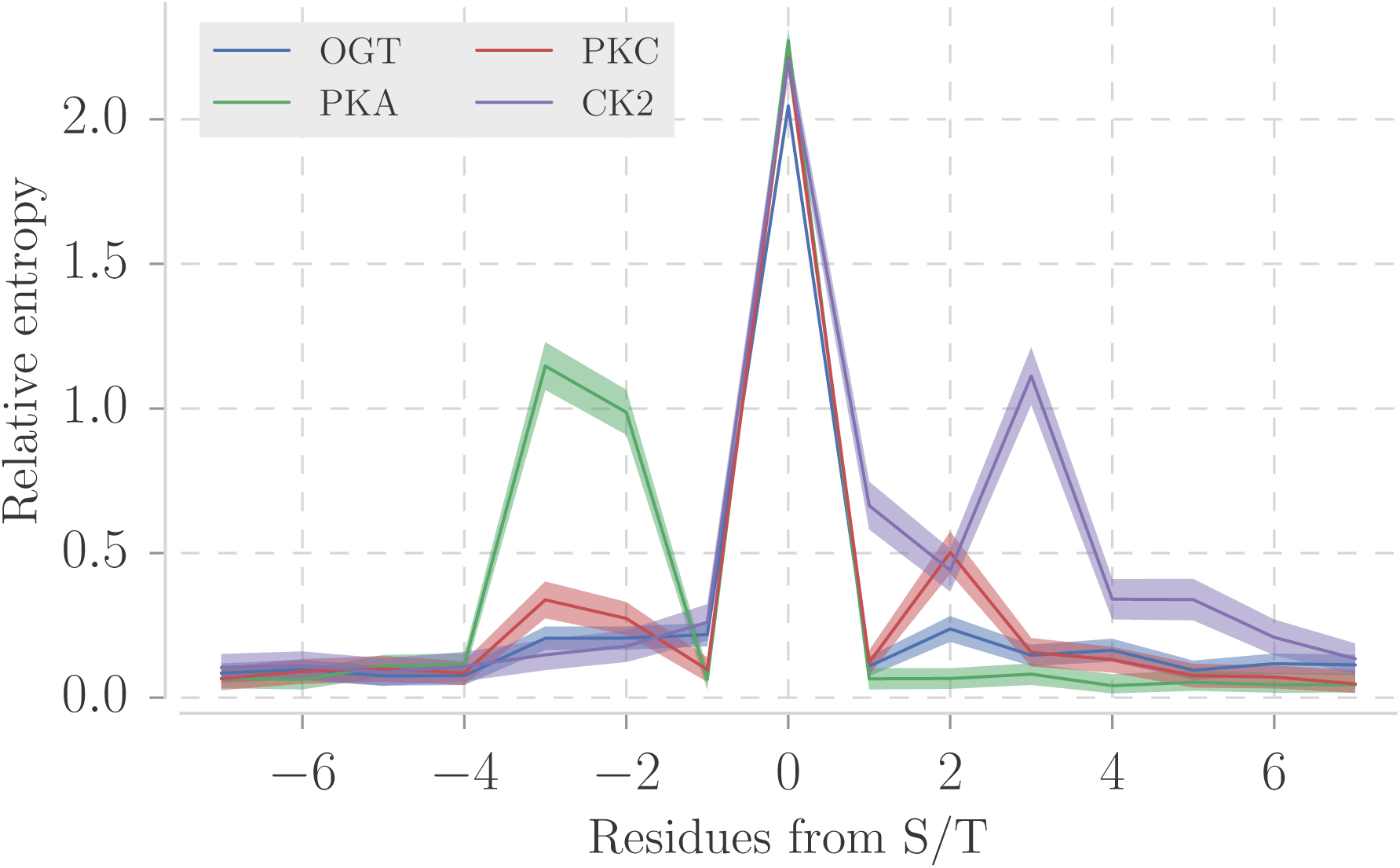
Sequence relative entropy of sites (+/- 7 residues) from 4 post-translational modifications. Three kinases with most sites in PhosphoSitePlus database^14^: protein kinase A (PKA with 1285 sites), protein kinase C (PKC with 930 sites) and casein kinase 2 (CK2 with 742 sites). 1530 OGT sites were compiled from the same database. The sequence relative entropy was calculated with the WebLogo library^43^. Lines show mean relative entropy and the semitransparent area represents 95% confidence intervals. Unlikely the PKA, PKC and CK2, known OGT sites have no clear sequence consensus.

The crystal structure of OGT in a ternary complex with UDP–GlcNAc and a peptide substrate revealed that the OGT and the peptides' residues predominantly make contact via the peptide backbone^21, 22^. This fact reduces the importance of the peptide side chain in the enzyme active site, the cleft where the reaction occurs. A short structural motif, instead of sequence motif, could work as a point of molecular recognition even with a degenerate sequence. Accordingly, in this paper, the three-dimensional structures of S/T OGT substrates were examined to determine if they have distinct structural motifs and patterns of secondary structure or solvent accessibility. In addition, the predicted secondary structure and disorder were compared for known OGT substrates and S/T unlikely to be modified.

## II. Methods

### A. Data sources

The data selection process is summarised in Figure 2. A total of 1,533 modified sites from 676 proteins were selected by combining proteins curated from the literature up until 2011^18^ and from 2011-2013^20^. The sites were filtered to keep 7-residue long motifs with unique sequences. The resulting dataset contained 1,385 sites in 620 proteins. This dataset is referred to hereafter as the “modified sequence sites” (MSS). For comparison, 100,329 S/T from the same proteins, but not annotated as OGT-modified, were selected as a background and are referred to here as the “unmodified sequence sites” (USS).

**Figure 2:**
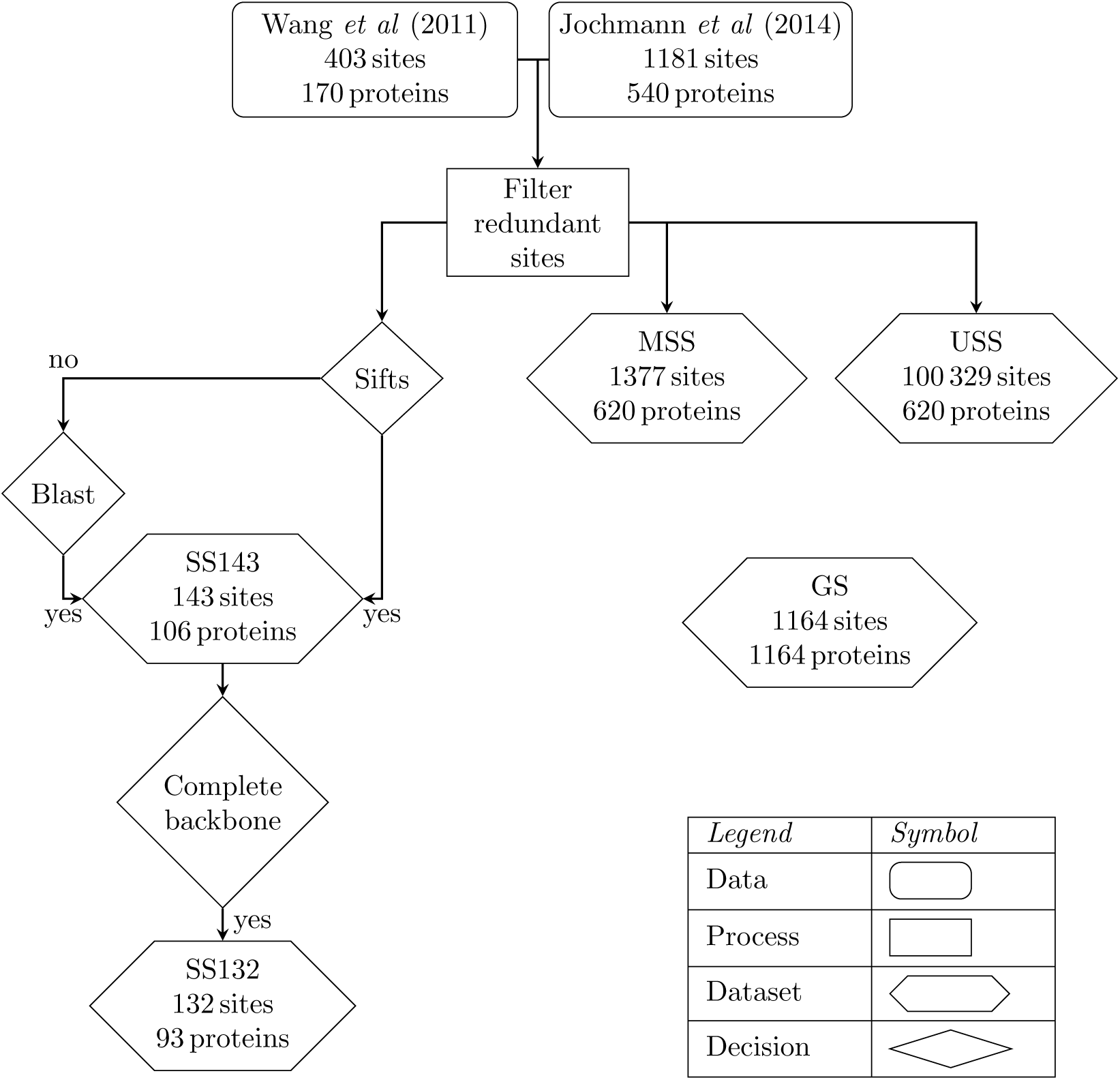
Diagram of the relationships among the 5 datasets used in this work. See Methods for details.

### B. Mapping O-GlcNAc sites to protein structures

Protein chains > 30 residues long from structures determined by X-ray crystallography to ≤ 2.50 Å resolution were selected from the Protein Data Bank^23^ (PDB: 2^nd^ August of 2015). Mapping the 1,385 OGT sites from 620 proteins to PDB structures by SIFTS^24^ located 45 OGT sites in 24 proteins of known structure. The structures of a further 107 sites were identified by searching the sequences of O-GlcNAcylated proteins against the PDB chains with BLAST and filtering by a conservative E-value ≤ 10^−25^ to minimise erroneous matches. The cutoff of ≤ 10^−25^ was found empirically to ensure the reliability of the match in the region of each site by inspecting all alignments between query and PDB sequence at different thresholds. Selecting the protein chain with highest coverage (SIFTS) or E-value (BLAST) left 143 sites in 107 proteins for further analysis, referred to hereafter as the “143 Structural Sites” (SS143).

### C. Site definition and clustering

The three-dimensional structure of OGT with its substrates suggests the region of contact between OGT and a modifiable S/T includes the residues and +/- 3 amino acids either side^21, 22^. From the structural sites returned in Mapping O-GlcNAc-sites to protein structures, “132 Structural Sites” (hereafter SS132) had at least one match with all backbone atoms for the 7-residue long site and were retained for further analysis. Cα atoms of each residue and the Cα and the Cβ for the central S/T were superimposed for all pairs of sites. Hierarchical clustering by complete linkage was applied on the resulting matrix of root-mean-square deviation (RMSD) values and clusters selected where all pairs of peptides were within 3 Å RMSD of each other.

### D. Structural properties of sites

Protein secondary structure assignments were obtained from DSSP^25^. DSSP annotates 7 different secondary structure states: 3_10_ helix (G), α helix (H), π helix (I), bends (S), turns (T), isolated (B) and extended (E) β-bridge. These assignments were reduced to three states: G and H to helices (H); I, B and E to strands (E); and all other, including residues with no assignment, to coils (C)^26^. The solvent accessible area from DSSP was normalised by the residue's maximum accessible area^27^. A S/T was considered exposed if its relative solvent accessibility (RSA) was > 25%; partially buried if the RSA > 5% and ≤ 25%, and buried if RSA ≤ 5%. Cα B-factors were standardised (Z-score normalised) over the B-factors for all Cα in the same chain.

### E. Prediction of protein disorder and secondary structure

Protein secondary structure predictions for the proteins in the MSS dataset were performed by JPred4^28^. Since JPred4 limits sequence longer than 800 residues, 300 of the sequences in the MSS dataset sequences were trimmed while ensuring the modified S/T was at least 100 residues away from the N- and C-termini to avoid edge effects. The intrinsic disorder was predicted by JRonn (Java implementation of Ronn^29^), IUPred^30^ and DisEMBL^31^ through the JABAWS^32^ command line application. Between them, these methods provide 6 different disorder prediction scores: DisEMBL-REM465 (0.6), DisEMBL-COILS (0.516), DisEMBL-HOTLOOPS (0.1204), IUPred-Long (0.5), IUPred-Short (0.5) and JRonn (0.5). The score ordered/disordered classes were defined by the cut-offs (in parenthesis) defined by the methods’ authors. Disorder predictions were also performed on a background set of 1,164 S/T selected at random from globular proteins in the Astral dataset^33^ version 2.04, referred to hereafter as the “Globular Set” (GS).

### F. Statistical analysis and code

The data collection, processing, analysis and the Cα clustering steps, were written in the Python programming language (Python Software Foundation, version 2.7 http://www.python.org) with the libraries Pandas (version 0.17)^34^ and Biopython (version 1.65)^35^. Statistical tests were performed with the StatsModels (version 0.6) and Scipy (version 0.16) libraries. A *p* value *(p*) threshold was set to 0.05.

## III. Results and Discussion

### A. Analysis of O-GlcNAc sites in proteins of known three-dimensional structure

Previous reports have suggested that O-GlcNAc sites, like phosphorylation sites, are predominantly present in disordered regions of proteins^36^. One indication of structural disorder is the crystallographic B-factor which indicates regions of the protein that lack crystallographic contacts. However, the standardised B-factor distribution on the SS143 dataset is the same for modified and unmodified S/T (Kruskal-Wallis two-sample test *p* = 0.12).

In X-ray crystal structures, the REM465 residue annotation indicates residues that are missing from the protein structure model and has previously been used as an indicator of structural disorder^31^. Of the 143 S/T in the SS143 dataset, 26 are in regions of the protein structure labelled as REM465. In comparison, 553 of 4,811 unmodified S/T from the same protein structures are also found in REM465 regions. Accordingly, O-GlcNAcylated S/T in these proteins are 1.7 times more likely to be in REM465 regions (Fisher's exact test *p* = 0.02). This finding is consistent with O-GlcNAcylated S/T occurring more frequently in disordered or highly flexible regions.

Table 2 summarises the DSSP assigned secondary structure for the SS143 compared to the 4,811 unmodified S/T in the same proteins. The proportions of H, E and C are equivalent for the two groups implying that there is no preference in the secondary structure for modified S/T in this dataset.

**Table 1.** Dataset summary. See Methods for details.

| Dataset name | Number of sites | Number of proteins | Short name |
| --- | --- | --- | --- |
| Modified Sequence Sites | 1,385 | 620 | MSS |
| Unmodified Sequence Sites | 100,329 | 620 | USS |
| Structural Sites | 143 | 106 | SS143 |
| Structural Sites with backbone | 132 | 93 | SS132 |
| Globular Set | 1,164 | 1,164 | GS |

**Table 2.** DSSP assigned secondary structure proportion of S/T in the SS143 dataset compared to unmodified S/T in same protein chains. 95% CI – 95% confidence interval; n - number of S/T. The *p* value refers to the two-tailed z-score test between the proportions of modified and unmodified groups.

| Secondary structure | Modified |  | Unmodified |  | *p* value |
| --- | --- | --- | --- | --- | --- |
|  | Proportion (n) | 95% CI [lower, upper] | Proportion (n) | 95% CI [lower, upper] |  |
| C | 0.55 (78) | [0.46, 0.63] | 0.51 (2475) | [0.50, 0.53] | 0.36 |
| H | 0.25 (36) | [0.18, 0.32] | 0.32 (1525) | [0.31, 0.33] | 0.06 |
| E | 0.20 (29) | [0.13, 0.27] | 0.17 (811) | [0.16, 0.18] | 0.27 |
| Total | 143 |  | 4,811 |  |  |

Residues that are buried in the protein structure are not thought to be targeted by protein kinases, due to structural constraints. Figure 3 illustrates that there is no difference between modified and unmodified S/T with respect to relative solvent accessibility (RSA). 45% (65) of S/T in the O-GlcNAcylated proteins are exposed to solvent (RSA > 25%). Surprisingly, 7 O-GlcNAc sites, listed in Table 2, have an RSA < 5%, suggesting they are inaccessible to OGT in the natively folded protein.

**Figure 3:**
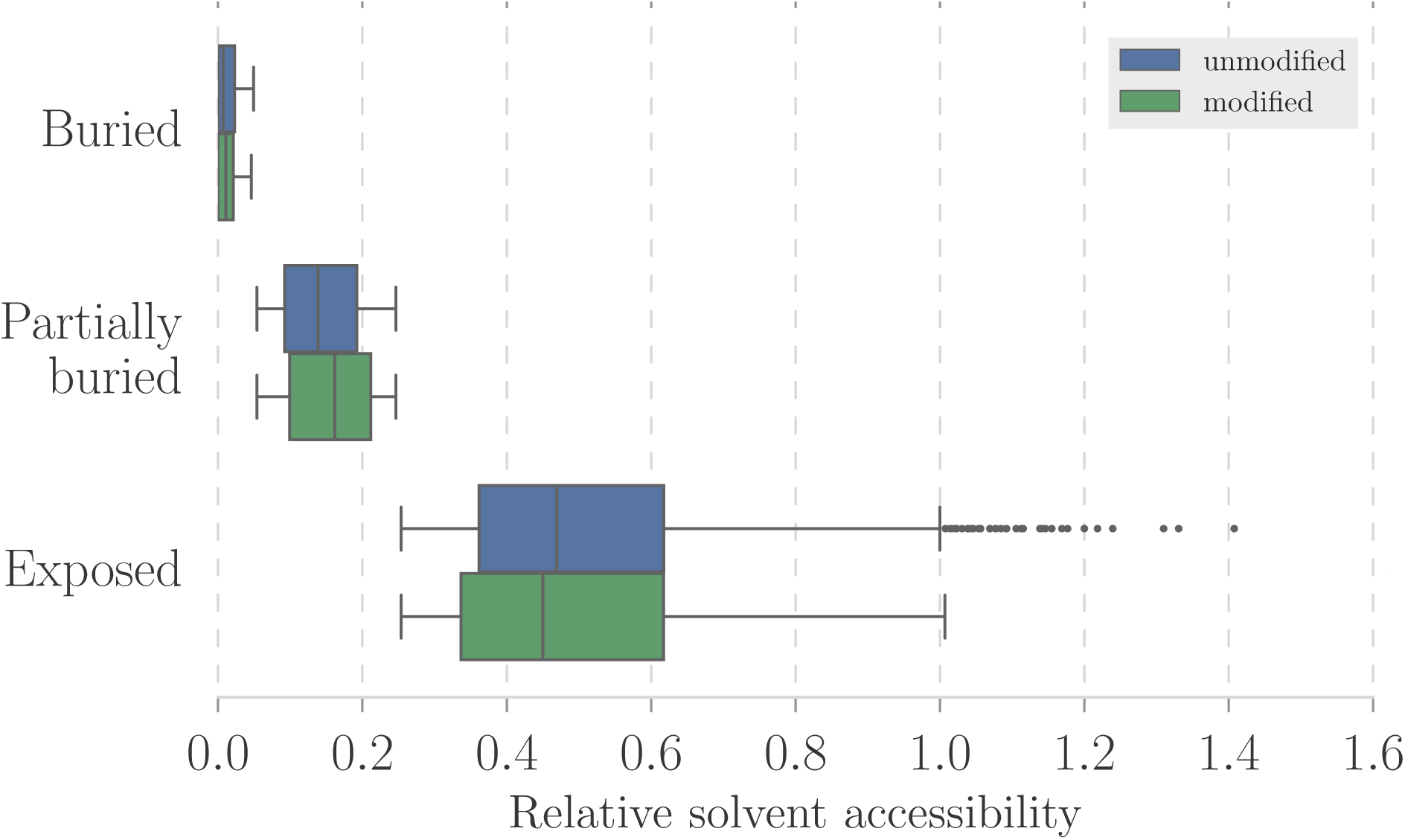
RSA of modified S/T in the SS143 dataset and unmodified S/T in same proteins. DSSP calculated solvent accessibility was normalised by the residue theoretical maximum accessibility, and the derived scores were reduced to three levels: buried (RSA ≤ 0.05), partially buried (0.05 < RSA ≤ 0.25) and exposed (RSA > 0.25). The y-axis and x-axis carry the RSA levels and the RSA distribution for each level, respectively. The mean RSA is equivalent between modified and unmodified residues, at all three levels.

### B. Groups of sites with similar local structure

Since the secondary structure and relative accessibility of modified S/T were indistinguishable from unmodified S/T, the local structure of the 7 residue peptides centred on S/T was investigated by pairwise superposition and clustering (see Methods). 36 sites produce singlet clusters, while the remaining 96 sites fall into 10 clusters. Sites in clusters had less than 3 Å RMSD from each other. Figure 4 illustrates the superimposed structures for sites in clusters, where green, yellow and grey represent residues in H, E, C secondary structures, respectively. The clusters show that sites are found in a wide range of secondary structure states as summarised in Appendix 1. The sites in Clusters E, G and J, have consistent consensus secondary structures. Clusters A–D, F, H and I are all variants on coil-helix or coil-strand transitions.

The buried sites, which are listed in Table 3, group in clusters D and G. The 3 sites in cluster D are unlikely to be targeted by OGT because they are buried in the protein core. In contrast, the 2/4 sites in cluster G (structures 3abm and 4y7y) might be modified since are located at a dimer interface, and so the monomer could be modified. The remaining two sites in cluster G (structures 2zxe and 4l3j) lie on a loop that could potentially move to expose them to OGT.

**Table 3.** Structural evidence of buried *O*-GlcNAc sites in the SS143 dataset. RSA – site mean relative solvent accessibility; Cluster id – Clusters in Figure 4.

| PDB id | Chain | Position | Cluster id | RSA |
| --- | --- | --- | --- | --- |
| 1f4j | B | 114 | D | 0.05 |
| 3cb2 | B | 170 | D | 0.02 |
| 4qvp | T | 131 | D | 0.01 |
| 2zxe | A | 366 | G | 0.02 |
| 3abm | R | 63 | G | 0.01 |
| 4l3j | A | 180 | G | 0.01 |
| 4y7y | Z | 190 | G | 0.04 |

**Figure 4:**
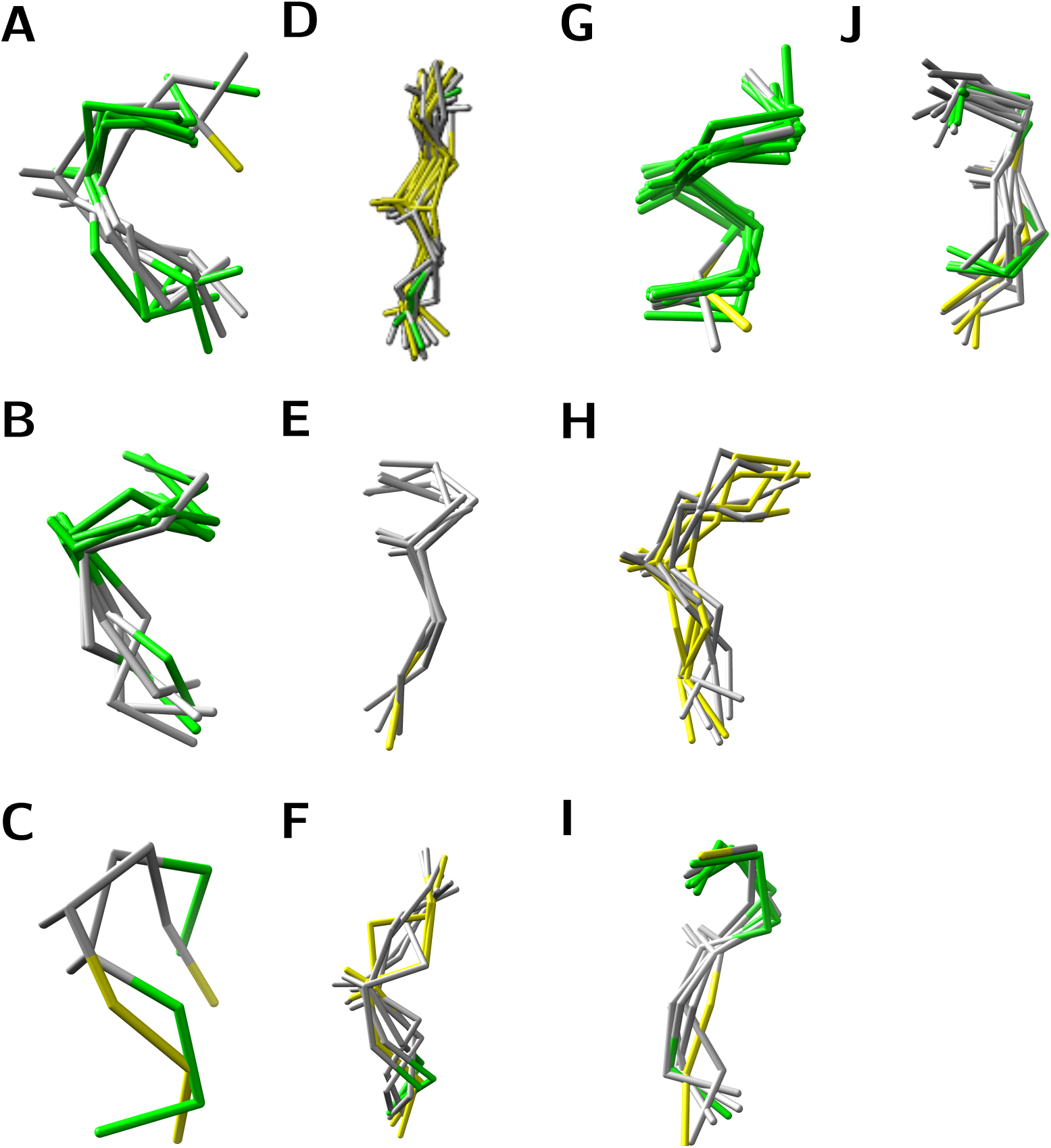
Structural superimpositions for the 10 clusters comprising 96 sites in the SS132 dataset. Pairs of sites were superimposed on their 7 Cα atoms and the Cβ of the central S/T. Their pairwise RMSD were clustered with complete linkage and Euclidean distance. Clusters were defined by a 3 A RMSD threshold. Green, yellow and grey represent residues in H, E, C secondary structures elements, respectively.

To see if the clusters found for the SS132 dataset are features of *O*-GlcNAc modification or just reflect the composition of the protein structures, 132 sites, centred on unmodified S/T, were randomly sampled with replacement from the same proteins and clustered. The process was repeated 1,000 times and the resulting clusters compared to those clusters in the SS132 dataset. The number of clusters identified in each sample ranged from 10-14 (95% CI), which is consistent with the SS132 dataset. Furthermore, the structural clusters identified for the random sampling included structural clusters similar to those for the modified sites, suggesting there are no dominant secondary structural or conformational patterns indicative of *O*-GlcNAc modified sites in the SS132 dataset.

### C. Analysis of features predicted for the “Modified Sequence Sites” dataset (MSS)

Since the structural analysis of *O*-GlcNAc sites is limited by the number of sites in proteins of known three-dimensional structure, prediction algorithms were applied to the sequences in the MSS and USS datasets, as detailed in Methods. The proportions of S/T in the levels of solvent accessibility predicted by JPred are equivalent in the MSS and USS datasets, as shown in Table 4. 1% of the S/T are predicted to be buried in the MSS and USS datasets. Again, the result is unexpected, since sites modified by PTM are thought to be accessible in the protein native fold.

**Table 4.** JPred4 predicted solvent accessibility for S/T in the MSS and USS datasets. The proportions of buried S/T as predicted by the Jnetsol method in JPred4. The proportions of buried S/T are significantly smaller for modified group. 95% CI – 95% confidence interval; n – number of S/T predicted to be buried. The *p* value refers to the two-tailed z-score test between the modified and unmodified groups.

| Buried at | Modified (MSS) |  | Unmodified (USS) |  | *p* value |
| --- | --- | --- | --- | --- | --- |
|  | Proportion (n) | 95% CI [lower, upper] | Proportion (n) | 95% CI [lower, upper] |  |
| 0% | 0.01 (7) | [0.00, 0.01] | 0.01 (836) | [0.008, 0.009] | 0.18 |
| 5% | 0.04 (55) | [0.03, 0.05] | 0.04 (3,917) | [0.038, 0.040] | 0.86 |
| 25% | 0.29 (403) | [0.27, 0.31] | 0.35 (28,044) | [0.27, 0.28] | 0.31 |

While the structural sites in the SS143 dataset have equal proportions of the secondary structure states, the result from secondary structure predictions on the MSS set showed that *O*-GlcNAc sites are likely to reside in coils, if compared to the USS dataset.

Table 5 shows an increase of the proportion of modified S/T in C (*p* < 0.01) and a corresponding reduction in H (p < 0.01), but no change in E (*p* = 0.6). The enrichment of sites in C is consistent with the need to place modified S/T in loops that are more likely to be mobile and so more accessible to OGT. The proportions of secondary structure assigned by DSSP and predicted by JPred4 differ. While secondary structure prediction has limited accuracy, the number of samples in the SS143 dataset is limited and potentially biased toward structured regions in proteins. Also, clustering sites in the SS132 dataset highlight groups that are more likely to occur near to the transition between a secondary structure element and C, as observed in several members of clusters A–D, F and H. The regions of transition between C and H/E are harder to predict than contiguous secondary structure elements, and this may also contribute to the observed enrichment in C.

**Table 5.** JPred4 predicted secondary structure proportions for S/T in the MSS 0025; confidence interval; n – the number of S/T; the p value refers to the two-tailed z sand USS datasets. 95% CI – 95%core test between the modified and unmodified groups.

| Secondary structure | Modified (MSS) |  | Unmodified (USS) |  | *p* value |
| --- | --- | --- | --- | --- | --- |
|  | Proportion (n) | 95% CI [lower, upper] | Proportion (n) | 95% CI [lower, upper] |  |
| C | 0.88 (1,205) | [0.86, 0.90] | 0.829 (83,150) | [0.826, 0.831] | <0.01 |
| H | 0.08 (106) | [0.07, 0.09] | 0.126 (12,684) | [0.124, 0.128] | <0.01 |
| E | 0.05 (66) | [0.04, 0.06] | 0.045 (4,495) | [0.044, 0.046] | 0.6 |

The analysis of SS143 dataset showed an enrichment of S/T in REM465 regions likely to be disordered or highly mobile. To explore this further, 3 disorder prediction algorithms, giving a total of 6 disorder scores, were run on the MSS and USS datasets as detailed in Methods. Table 6 shows that, with the exception of DisEMBL-HOTLOOPS which is trained structural B-factors, all methods report a small but significant increase in mean predicted disorder for the modified S/T. To confirm this result, the MSS dataset was compared to the GS dataset, which was selected from proteins known to be predominantly globular, and hence an ordered background. In Figure 5, DisEMBL-HOTLOOPS shows an increase in the ratio of disordered residues around the modified S/T. DisEMBL-COILS and JRonn also indicate a small increase, not in a specific region, but rather for 40 residues around the S/T. IUpred-Long, IUPred-Short and DisEMBL-REM465 show a bigger increase of the ratio of disordered residues in the MSS dataset and IUpred-Short and REM465 have a clearer peak within −15 to 15 residues from the modified S/T. Overall, all methods indicate an increased proportion of predicted disorder in the MSS dataset when compared to the GS dataset.

**Figure 5:**
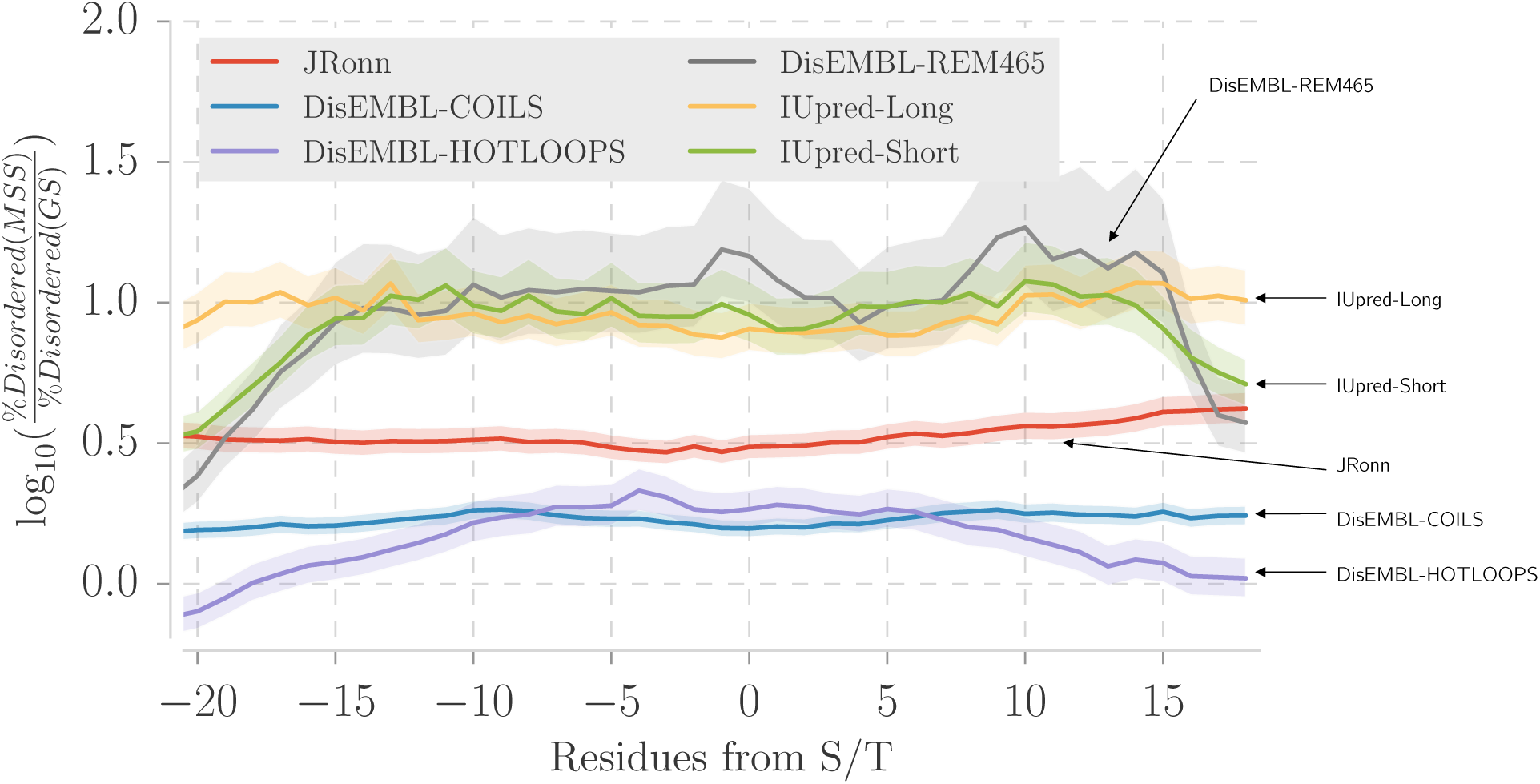
Predicted disorder around *O*-GlcNAc sites in the MSS compared randomly selected S/T in the GS-dataset. The y-axis shows the log_10_ odds ratio of the between the proportion of disordered residues in the MSS dataset and the proportion of disordered residues in the GS dataset. The semi-transparent area represents 95% confidence intervals. A residue was defined as disordered given a method threshold. The x-axis represents the distance in residues of the central residue, always a S/T. DisEMBL-REM465, IUpred-short predict protein structural disorder specifically around the modification site, while the other methods detect intrinsic disorder over O-GlcNAcylated proteins. DisEMBL-HOTLOOPS shows a less pronounced increase compared to the other methods.

**Table 6.** Predicted disorder between modified and unmodified S/T. All disorder prediction methods, excepting DisEMBL-HOTLOOPS, reveal a small but significant increase of mean disorder score for modified S/T over unmodified ones. The p value refers to the two-tailed t-test between the modified and unmodified groups. SE – standard error.

| Method | Mean score modified (MSS) ± SE | Mean score unmodified (USS) ± SE | *p* value |
| --- | --- | --- | --- |
| DisEMBL-REM465 | 0.48 ± 0.004 | 0.47 ± 0.001 | 0.01 |
| DisEMBL-COILS | 0.60 ± 0.004 | 0.58 ± 0.001 | <0.01 |
| DisEMBL-HOTLOOPS | $0.10 \pm 0.001$ | $0.10 \pm 0.001$ | 0.45 |
| IUpred-Long | $0.59 \pm 0.006$ | $0.55 \pm 0.001$ | <0.01 |
| IUpred-Short | $0.48 \pm 0.005$ | $0.45 \pm 0.001$ | <0.01 |
| JRonn | $0.62 \pm 0.004$ | $0.61 \pm 0.001$ | 0.02 |

## IV. Conclusions and final remarks

Despite the substantial evidence of protein structural disorder in the MSS and the SS143 datasets, the SS132 dataset clearly indicates that some of the examined sites appear within ordered regions of the protein structure. Furthermore, InterproScan^37^ analysis of *O*-GlcNAc sites assigned 19 *%* of the sites to protein domains, this is similar to with the 25 *%* phosphoserines and phosphothreonines in PFAM domains^14, 38^, which are thought to be mostly ordered by definition. So, like protein phosphorylation, *O*-GlcNAcylated S/T are found in both ordered and disordered regions.

The local tertiary structure of *O*-GlcNAc sites is indistinguishable from unmodified sites. So, how does OGT recognise the site it modifies? OGT may force the unfolding of the targeted substrate^39^. Moreover, OGT participates in macromolecular assemblies^40^, and so adaptor proteins should be important. Non-local interactions might also act in OGT substrate recognition. In protein kinase C (PKC) substrate recognition, residues distant in the protein sequence but close in its three-dimensional structure are critical^41^. Other components, such as UDP–GlcNAc concentration and subcellular location-dependent interactions, modulate OGT activity^42^, but their part in substrate recognition is still unknown. In conclusion, although no three-dimensional fingerprint was detected during the structural characterisation of OGT-modified sites, the work confirmed that S/T and surrounding residues are more disordered than the backgrounds tested and that sites in transition between C to H/E might be involved, suggesting that the structural flexibility has a role on OGT site recognition.

## Acknowledgements

We would like to thank Dr. Tom Walsh and the University of Dundee IT department for computing support; Prof. Daan van Aalten and DVA group for advice and discussions.

